# Antibody effector functions are required for broad and potent protection of neonates from herpes simplex virus infection

**DOI:** 10.1101/2023.08.29.555423

**Authors:** Matthew D. Slein, Iara M. Backes, Callaghan R. Garland, Natasha S. Kelkar, David A. Leib, Margaret E. Ackerman

**Affiliations:** Department of Microbiology and Immunology, Geisel School of Medicine at Dartmouth, Lebanon, NH 03756, USA; Thayer School of Engineering, Dartmouth College, Hanover, NH 03755, USA

**Keywords:** antibody engineering, Fc effector functions, neonatal HSV infections, neutralizing antibody, monoclonal antibody therapeutics

## Abstract

The failure of multiple herpes simplex virus (HSV) vaccine candidates that induce neutralizing antibody responses raises the hypothesis that other activities, such as Fc domain-dependent effector functions, may be critical for protection. While neonatal HSV (nHSV) infection result in mortality and lifelong neurological morbidity in humans, it is uncommon among neonates with a seropositive birthing parent, suggesting the potential efficacy of antibody-based therapeutics to protect neonates. We therefore investigated the mechanisms of monoclonal antibody (mAb)-mediated protection in a mouse model of nHSV infection. Both neutralization and effector functions contributed to robust protection against nHSV-1. In contrast, effector functions alone were sufficient to protect against nHSV-2, exposing a functional dichotomy between virus types that is consistent with vaccine trial results. Together, these results emphasize that effector functions are crucial for optimal mAb-mediated protection, informing effective Ab and vaccine design, and demonstrating the potential of polyfunctional Abs as potent therapeutics for nHSV infections.

**Graphical abstract.**
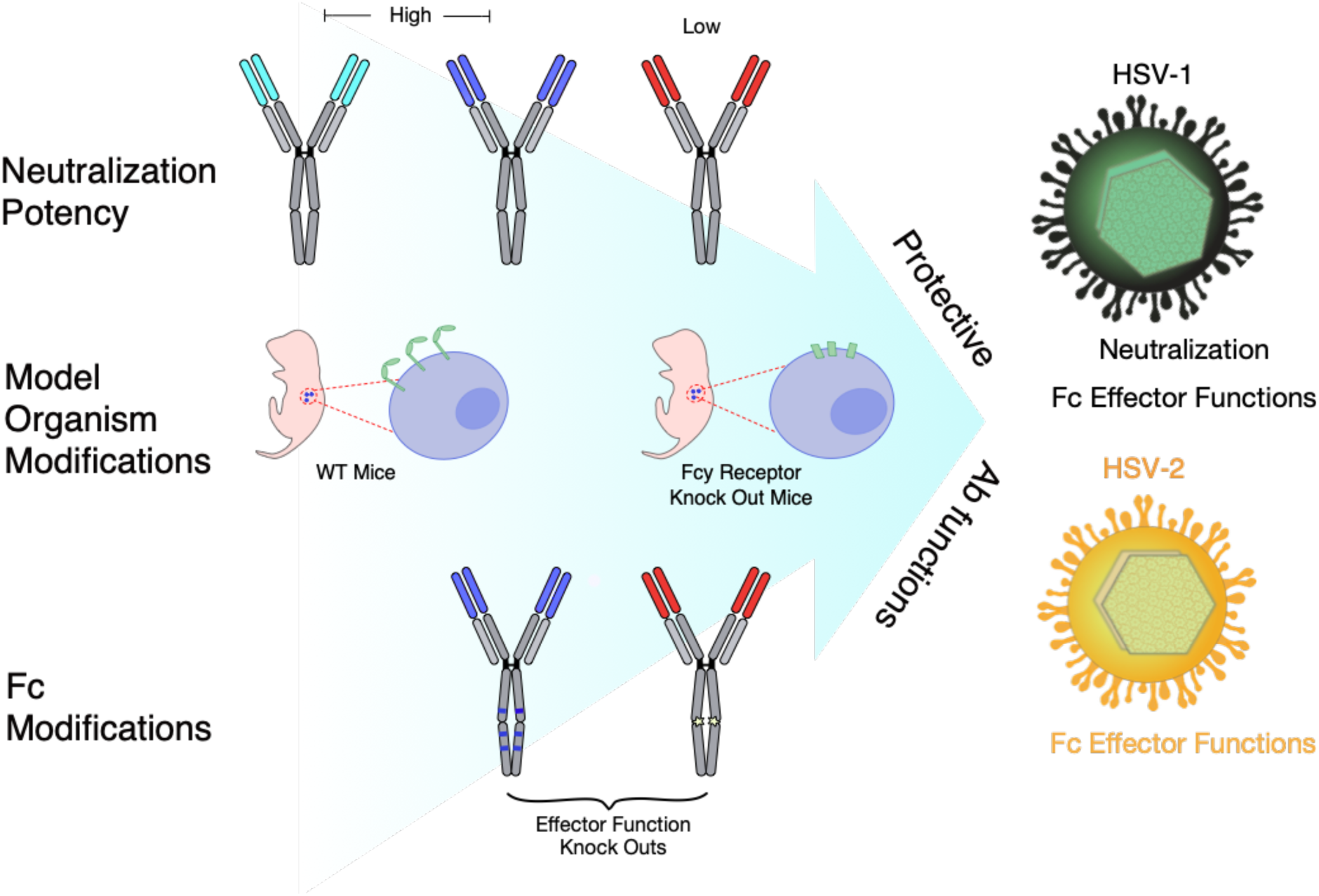
Mechanistic dissection of antibody-mediated protection from HSV. Monoclonal antibodies (mAbs) with varying neutralizing potencies and Fc modifications that impact effector function were evaluated in wildtype (WT) and FcγR-/- mice to define mechanisms of antibody-mediated protection from HSV infection. To model human vulnerability to HSV disease during the neonatal period, neonatal mice were challenged with HSV, treated with mAb, and then assessed for morbidity and mortality. We observed that polyfunctional mAbs provide broader and more potent protection than antibodies with either low neutralization or low effector function. Moreover, while sufficient for protection against HSV-1, neutralization activity alone was unable to protect from HSV-2 infection.

## Introduction

When encountered during the neonatal period, herpes simplex virus (HSV) infections can result in loss of life or long-term neurological disability^1–3^. Neonatal infections can present as skin, eye, and mouth (SEM) disease, which is amenable to antiviral therapy, or more invasive disseminated and/or central nervous system (CNS) disease. While new treatment regimens with acyclovir and its derivatives have improved outcomes, mortality following disseminated disease remains unacceptably high^4,5^. Most neonatal HSV (nHSV) infections are vertically transmitted during birth from a recently infected birthing parent who has not yet developed a mature antibody response to HSV type 1 or type 2 (HSV-1, HSV-2)^6^. Given the severity of neonatal infection resulting from primary maternal infection^6,7^, birthing parent seropositivity is believed to be protective due to the transfer of HSV-specific antibodies (Abs) via the placenta^2,5,8^. High titers of neutralizing or antibody-dependent cellular cytotoxicity (ADCC)-inducing Abs in infected neonates have been associated with less severe disease^9,10^ . Animal studies support the notion that neutralization and Fc-effector functions, such as ADCC, antibody-dependent cellular phagocytosis (ADCP), and antibody-dependent complement deposition (ADCD) can aid in the clearance of acute HSV infection^11–14^. Further insights into how antibodies exert direct and indirect antiviral activities to protect against infection could aid in the design of both passive and active immunization strategies for HSV.

To this end, whether neutralization or effector functions play a dominant role in protection from HSV-mediated disease has long been unclear, as conflicting results have been reported in animal models^12,13,15–17^. Previous studies differentiated effector functions from neutralization by treating with digested antibody (Fab) fragments^18^. However, digestion is known to compromise neutralization potency and half-life, which confounds interpretation of study results. Other studies have sought to answer this question using polyclonal Ab or mAbs that could either neutralize or carry out specific effector functions^10,19^. While such approaches have contributed to our understanding of the potential contributions of Ab effector functions, disparities in protection from disease could also be attributed to the specific epitope(s) targeted, differences in Ab affinity or avidity, or other factors. Ab Fc engineering strategies that allow separation of Fc-dependent effector functions from neutralization provide a platform to improve experimental resolution in defining Ab-dependent mechanisms of protection^20–23^, which can inform both vaccine design and therapeutic mAb development.

Like other consequential early life pathogens, most studies of HSV have focused on adult animal models. There is therefore a dearth of information on how Abs protect in the neonatal period. Given this knowledge gap, we sought to investigate the mechanism(s) by which Abs that target glycoprotein D (gD) mediate protection against nHSV-1 and nHSV-2 infections. Using a mouse model of nHSV infection, we demonstrate that there are distinct mechanisms of Ab-mediated protection that differ between viral types, motivating the optimization of Ab therapeutics that could ameliorate nHSV. Given the short time window of vulnerability to nHSV, this work could facilitate the design of effective therapeutic mAbs, whose timely administration could yield tremendous benefit for this devastating disease.

## Results

### Characterization of HSV-glycoprotein D (gD) specific monoclonal antibodies

The mAbs used in this study protect both adult and neonatal mice from HSV-1- and HSV-2-induced mortality^14,24–26^, but the mechanisms by which they elicit protection have not been defined. In order to better understand the contribution of neutralization and other Fc-mediated functions, we studied UB-621, HSV8, and CH42 AAA, mAbs which exhibit different neutralization potencies and effector function activity (**Figure 1A, Table S1**). To probe the contributions of effector functions *in vivo*, HSV8 and CH42 AAA were expressed with Fc domain point mutations that serve as functional FcγR and C1q binding knock outs (KO). UB-621 and HSV8 are unmodified human IgG1 mAbs, while CH42 AAA has been engineered with S298A/E333A/K334A mutations, which increase affinity for FcyRIIIA^27^. For construction of KO mAbs, we incorporated LALA PG^28^ mutations into HSV8 and the N297A^29^ substitution into CH42. VRC01^30^, an HIV-specific IgG1 mAb was included as an isotype control. The Fc receptor (FcR) binding profiles of the engineered mAbs were evaluated *in vitro* (**Figure 1B, Figure S1)**. The binding patterns of all three Fc-intact antibodies, UB-621, HSV8, and CH42 AAA were comparable, with CH42 AAA exhibiting the strongest binding to all human and mouse FcRs tested. As expected, the HSV8 LALA PG variant displayed diminished binding to both human and mouse FcRs as compared to HSV8. The CH42 NA variant also exhibited diminished binding to human and mouse FcRs, with the exception of murine FcyRI, to which binding was only slightly diminished. Importantly, given our use of these mAbs in mouse experiments, the Fc-modified and -unmodified forms of each HSV-specific mAb displayed comparable binding profiles to the four mouse FcR as to their human counterparts. These data indicate a high level of concordance between species.

**Figure 1:**
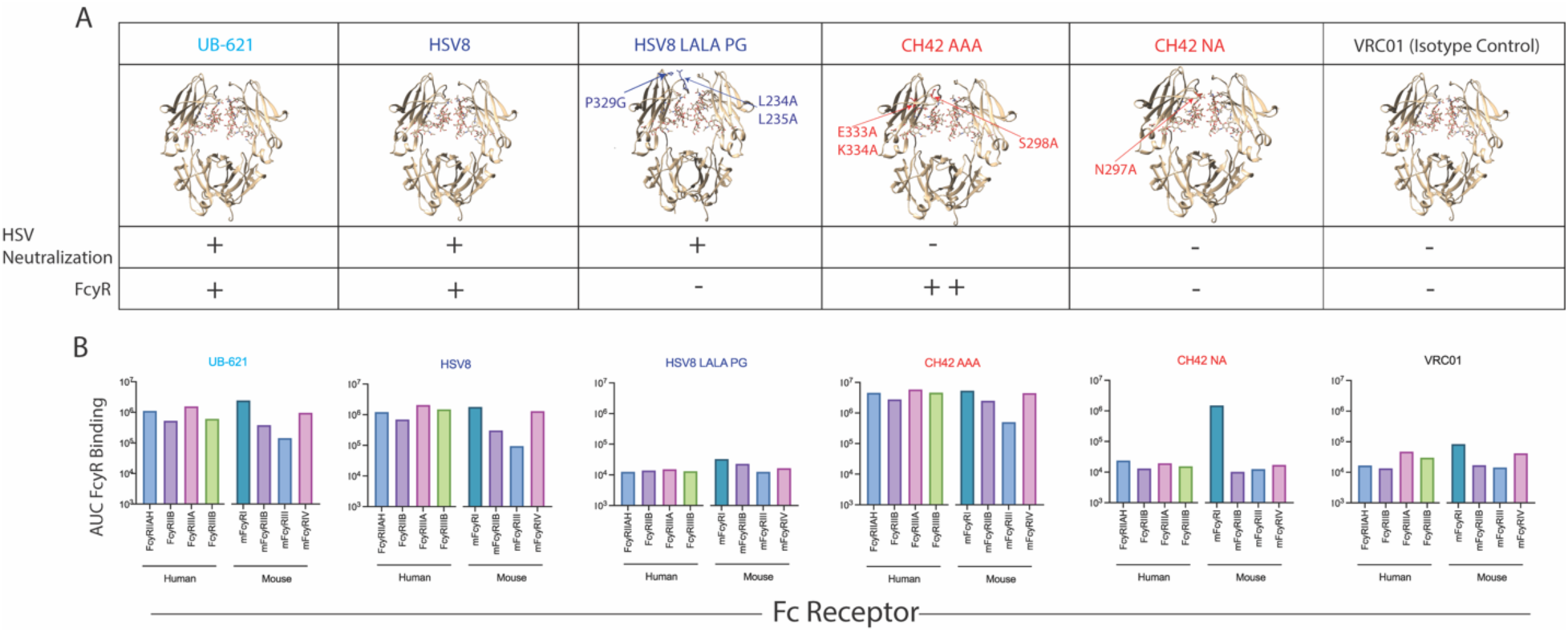
Biophysical Characterization of HSV gD-specific mAbs. **A**. Visualization of the Fc domains of mAbs used in this study. For reference purposes, mutated positions in the HSV-mAbs are superimposed on the crystal structure of the Fc domain of the HIV-specific mAb b12 (PDB: 1HZH). Reported neutralization potencies of each mAb and the expected ability of each Fc domain to bind FcyRs are indicated. **B**. FcyR binding profiles of the mAbs used in this study. Bar graphs present the area under the curve (AUC) for the binding of each mAb to recombinant human (left) and mouse (right) Fc receptors. Orthologous human and mouse Fc receptors are color matched.

To more directly assess the function of each mAb, *in vitro* assays of antigen recognition, neutralization, and effector function were performed (**Figure 2)**. Each HSV-specific mAb bound to both recombinantly expressed gD and gD expressed on the surface of mammalian cells (**Figure 2A-B, Figure S2B**). In contrast, the isotype control, VRC01, showed no binding. Notably, while CH42, HSV8, and UB-621 exhibited different antigen binding dose-response profiles from each other, the binding of Fc KO forms of HSV8 and CH42 to antigen were unchanged. Furthermore, direct antiviral activity afforded by antigen recognition again varied by mAb, but not by Fc modification (**Figure 2C-D**). UB-621 and HSV8 potently neutralized both HSV-1 and HSV-2, while CH42 poorly neutralized both viruses. Consistent binding and neutralization activities of unmodified and Fc KO mAbs permits the isolation of Fab- from Fc-dependent activities.

**Figure 2:**
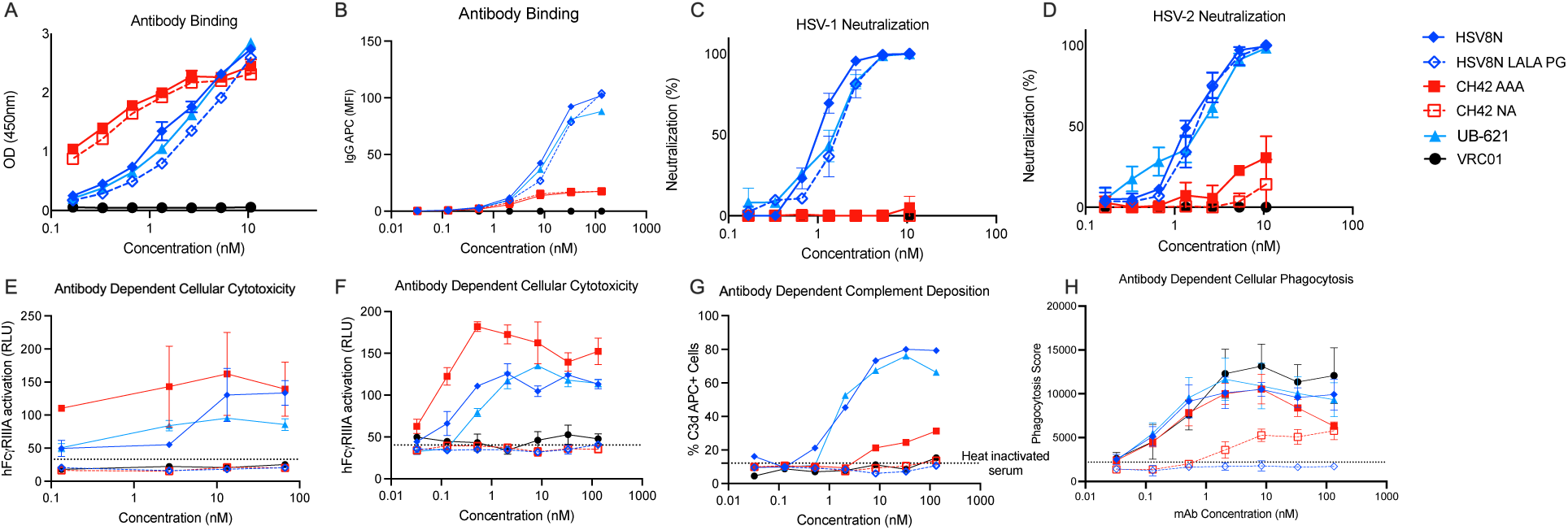
*In vitro* functional characterization of HSV gD-specific mAbs. **A-B**. Ability of the HSV gD mAbs to bind recombinant (**A**) or cell surface-expressed (**B**) gD via ELISA or flow cytometry, respectively. **C-D**. Ability of the HSV gD mAbs to neutralize HSV-1 (**C**) or HSV-2 (**D**) by plaque reduction assay. **E-H**. Effector function of HSV gD mAbs, including human FcγRIIIA stimulation of a reporter cell line in the context of antibody-bound gD on a microtiter plate (**E**), or gD-expressing cells (**F**) as surrogates for ADCC activity, complement deposition (**G**), or phagocytosis (**H**). Error bars represent standard deviation from the mean. OD – optical density, MFI – mean fluorescent intensity, RLU – relative light units, APC – Allophycocyanin.

Lastly, we tested the *in vitro* effector functions of these mAbs. We profiled their ability to promote FcyRIIIA activation upon recognition of recombinant or cell-expressed gD as a surrogate for ADCC activity (**Figure 2E-F**). We also measured their ability to induce complement deposition and phagocytosis (**Figure 2G-H, Figure S2A, C)**. HSV8, UB-621, and CH42 AAA all elicited effector functions *in vitro*, whereas KO mAbs were unable to elicit FcyRIIIA activation, complement deposition, or phagocytosis. As may have been anticipated from stronger binding to FcyRIIIA and FcyRIIA, CH42 AAA exhibited the most potent ADCC and phagocytic activity, indicating that the AAA mutations enhanced the ability of CH42 to elicit Fc function. CH42 NA, which eliminates the conserved Fc-glycan, did maintain some phagocytic activity, presumably due to residual binding to human FcyRI. Consistent with this observation, others have reported that aglycosylated IgG1 mAbs retain phagocytic activity via FcyRI expressed on macrophages^31,32^. Taken together, these experiments demonstrated the divergent activities of the mAb panel, supporting its utility to defining *in vivo* mechanisms of action.

### Neutralization and Fc-mediated functions contribute to nHSV survival

To begin to understand the roles of viral neutralization and Fc effector functions in mediating protection against a neonatal HSV challenge, 2-day old C57BL/6J pups were given intraperitoneal (i.p.) injections with 40 μg mAb, then immediately challenged with 1.0 x 10^4^ PFU HSV-1 intranasally (i.n.). Pups that received potently neutralizing mAbs HSV8 or UB-621 had improved survival as compared to pups that received non-neutralizing mAb (CH42 AAA) (**Figure 3A-B**). That said, all three HSV-specific antibodies improved survival as compared to isotype control (VRC01) (**Figure 3A-C**). HSV-infected mice treated with CH42 NA, which lacks both neutralization and effector function activity, succumbed to infection (**Figure 3B**). In contrast, the mice that received the neutralizing but effector function KO HSV8 LALA PG, survived HSV-1 infection (**Figure 3A**). These results indicate that for HSV8, neutralization alone was sufficient to mediate protection, while the moderate protection mediated by CH42 AAA was wholly Fc-dependent.

**Figure 3:**
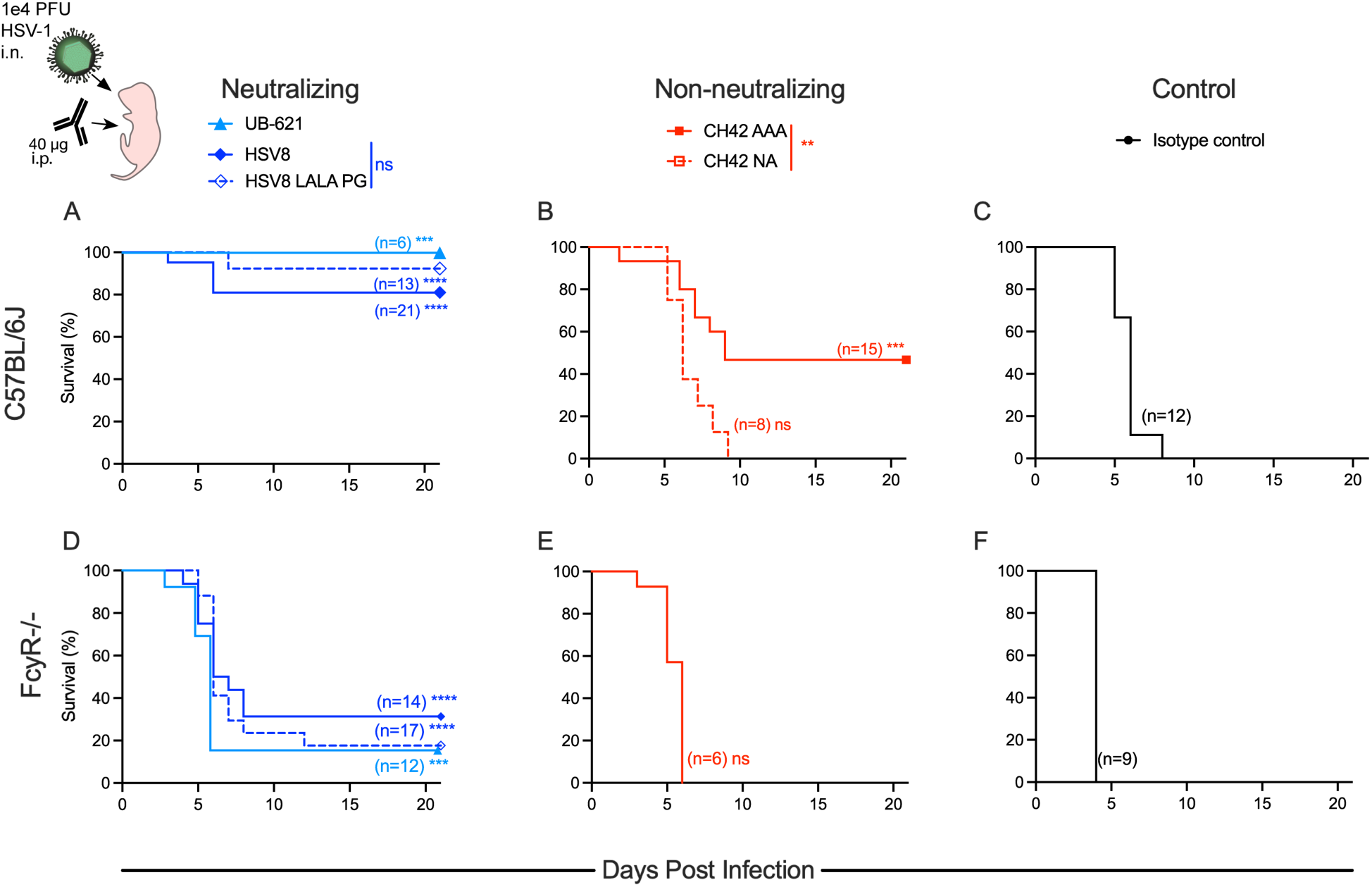
Both neutralization and effector function contribute to mAb-mediated protection from lethal HSV-1 challenge. Immediately before lethal intranasal (i.n.) challenge with 1×10^4^ plaque forming units (PFU) of HSV-1, two-day old pups were administered 40 μg of mAb by intraperitoneal (i.p.) injection. **A-C**. Survival of C57BL/6J pups receiving neutralizing mAbs UB-621, HSV8, or HSV8 LALAPG (**A**), non-neutralizing mAbs CH42 AAA or CH42 NA (**B**), or isotype control mAb VRC01 (**C**). **D-F**. Survival of FcyR-/- pups receiving neutralizing (**D**), non-neutralizing (**E**), or isotype control mAb (**F**). Number of mice in each condition and statistical significance as compared to isotype control in matched mouse strain determined by the Log-rank (Mantel-Cox) test (***p<0.001, ****p<0.0001) are reported in inset. Significance between HSV8 and HSV8 LALA PG or CH42 AAA and CH42 NA are reported in the above legend as determined by the Log-Rank (Mantel-Cox) test.

As an orthogonal test to define the specific contribution of FcyR-dependent Fc-effector functions in mediating protection, FcγR-deficient mice (FcyR-/-) ^33^ were treated i.p. with mAbs and challenged i.n. with 1.0 x 10^4^ PFU HSV-1. FcyR-/- mice that received neutralizing mAbs HSV8, HSV8 LALA PG, and UB-621 exhibited increased survival as compared to CH42 AAA and control IgG (**Figure 3D-F**). They were, however, considerably more susceptible to HSV infection as compared to C57BL/6J wild type mice. Viral neutralization was highly protective in WT mice but not in FcyR-/-mice, indicating a role for Fc function in contributing to protection against HSV-1 in neonatal mice.

### Fc-functions are protective in the absence of complete viral neutralization

Given the increased mortality observed in FcyR-/- mice treated with potently neutralizing mAbs, we next investigated the role of Fc-functions under conditions of maximal viral neutralization. Achieving maximal neutralization activity was accomplished by pre-incubating excess mAb with 1.0 x 10^4^ PFU of HSV-1 prior to *in vivo* challenge. With this experimental design, both C57BL/6J and FcγR-/- mice were completely protected from disease by HSV8 and UB-621 (**Figure 4A, D**). In contrast, when virus was pre-incubated with 20 µg of CH42 AAA, the majority of the pups succumbed to infection (**Figure 4B**), as did all animals treated with the isotype control mAb (**Figure 4C**). While increasing the CH42 AAA concentration fivefold to 100 µg of mAb/pup did improve survival (**Figure 4B**), it was unable to achieve the complete protection seen when mice were administered neutralizing mAbs. In contrast to neutralizing mAbs, even a 100 μg dose of CH42 AAA failed to protect FcγR-/- pups (**Figure 4E**), who exhibited survival comparable to the isotype control mAb (**Figure 4F**). These results provide evidence that FcγR-mediated activities are not necessary to provide protection in the context of fully neutralized HSV-1 but can be responsible for protection in the absence of complete neutralization.

**Figure 4:**
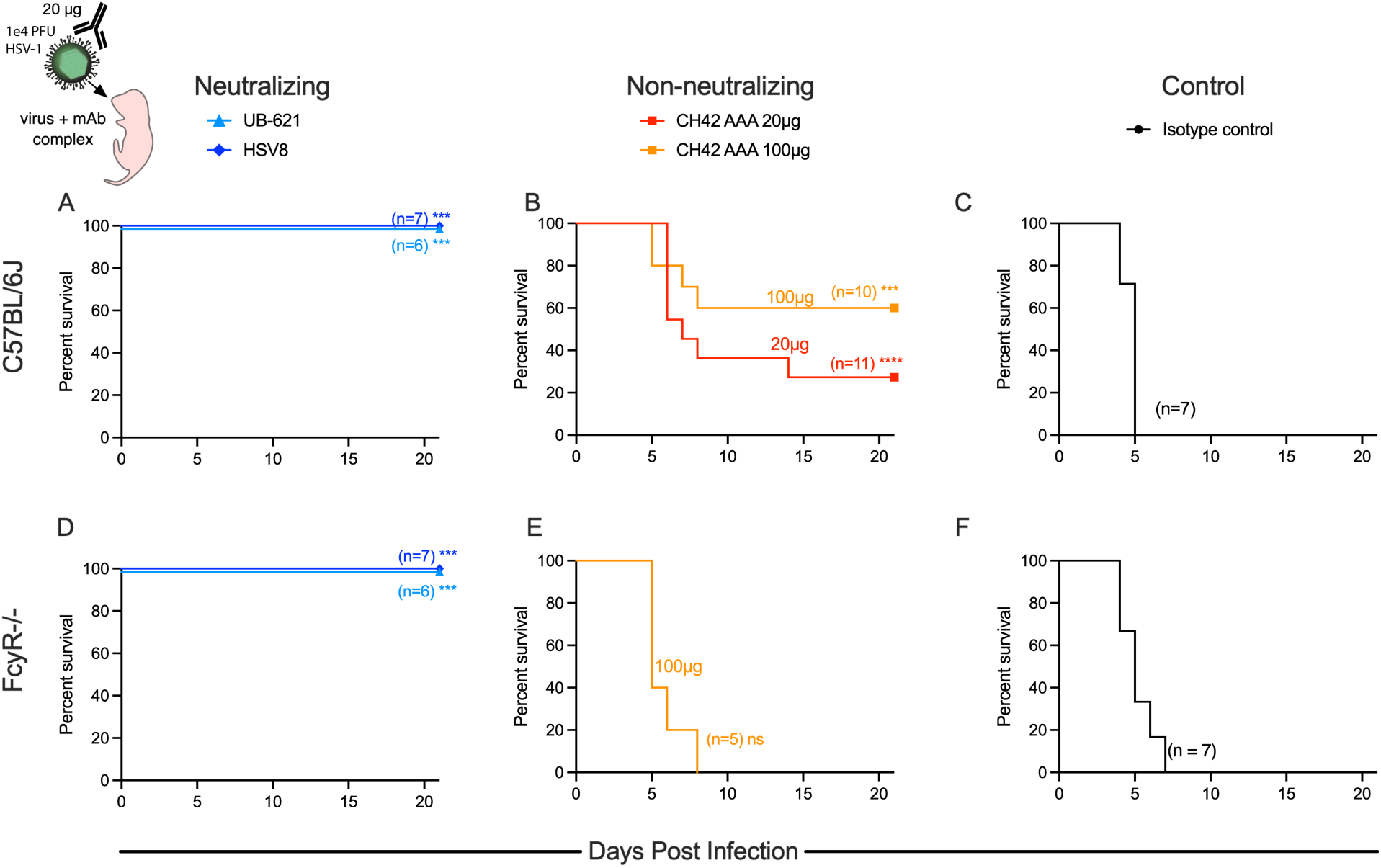
Fc functions are protective in the absence of complete viral neutralization. One hour before i.n. challenge of two-day old pups, immune complexes were formed by incubation of 1×10^4^ plaque forming units (PFU) of HSV-1 with mAb at 37°C (20 μg unless otherwise noted). **A-C**. Survival of C57BL/6J pups following immune complex challenge with virus opsonized with neutralizing mAbs UB-621 or HSV8 (**A**), non-neutralizing mAb CH42 AAA (20 µg or 100 µg) (**B**), or isotype control mAb (**C**). **D-F**. Survival of FcyR-/- pups following immune complex challenge with virus opsonized with neutralizing mAbs UB-621 or HSV8 (**D**), non-neutralizing mAb CH42 AAA (100 µg) (**E**), and isotype control mAb (**F**). Number of mice in each condition and statistical significance as compared to isotype control in matched mouse strain determined by the Log-rank (Mantel-Cox) test (**p < 0.01, ***p < 0.001, ****p < 0.0001) are reported in inset.

### The relative impacts of neutralization and Fc-effector functions are mAb dose-dependent

There was no difference in survival of pups treated with HSV8 or HSV8 LALA PG mAbs at the 40 µg dose, and full protection of FcγR-/- mice with neutralizing mAb-opsonized virus was observed. Together, these data indicate the lack of a major role for Fc-effector functions in mediating protection in the context of high levels of neutralizing activity. We wished, therefore, to assess the hypothesis that Fc-effector functions may be more important at lower antibody doses^34^. To test this possibility, we treated C57BL/6J mice with 10 µg mAb delivered i.p. and subsequently challenged with a lethal dose of HSV-1. As expected, this dose of HSV8 was less protective than the 40 µg dose. However, this lower dose of HSV8 LALA PG was completely unable to protect (**Figure 5A**), demonstrating that neutralization alone is insufficient to protect mice at lower mAb doses. Intriguingly, CH42 AAA provided comparable protection to HSV8 at the 10 µg dose (**Figure 5B**). In contrast, 10 µg of CH42 NA and the isotype control failed to protect mice from HSV-mediated mortality (**Figure 5C**). Effector functions, therefore, mediate protection from HSV-1-induced mortality at low antibody concentrations, at which viral neutralization may be incomplete.

**Figure 5:**
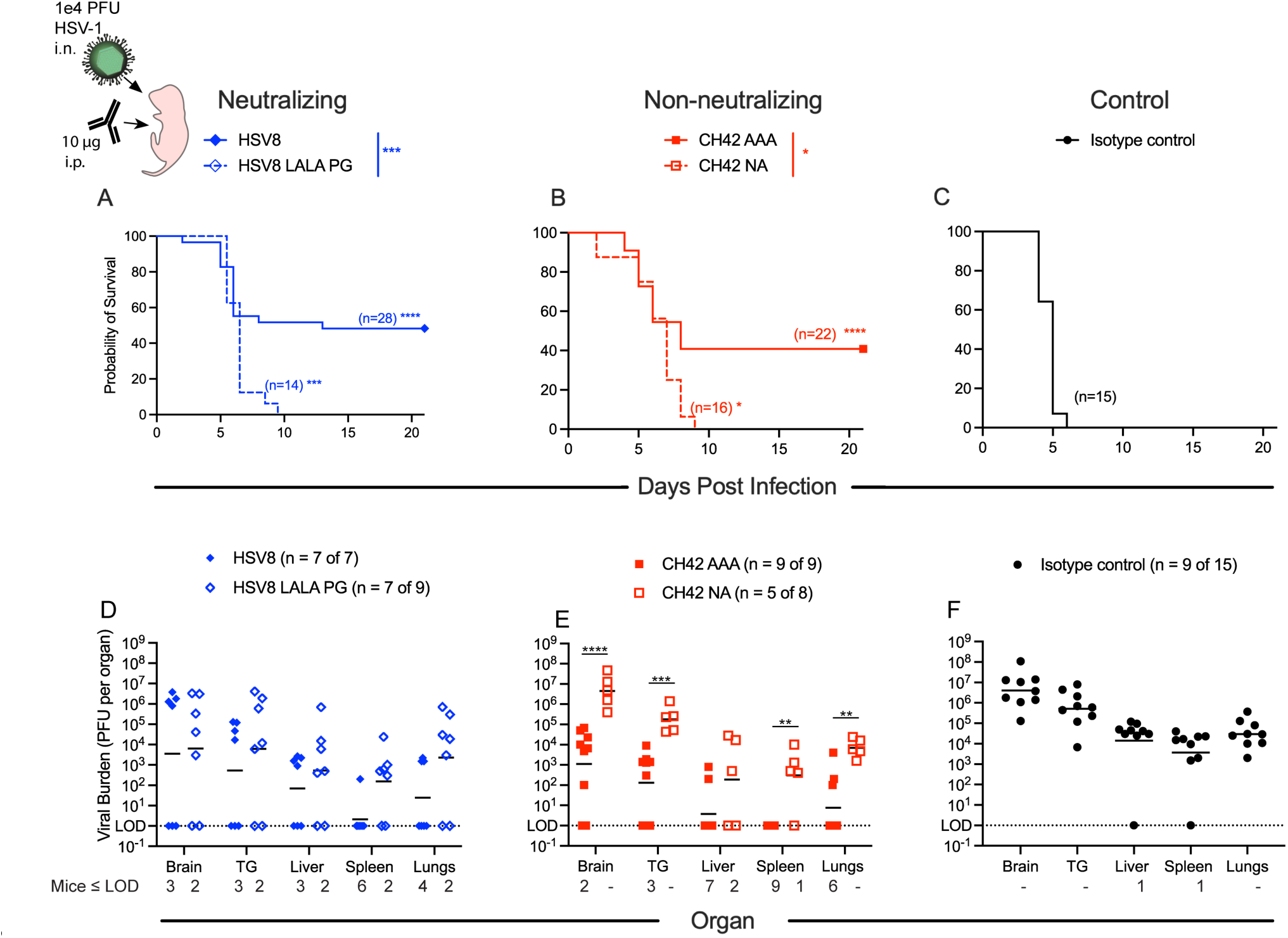
Relative contributions of neutralization and effector functions to protection from lethal challenge depend on antibody dose. Immediately before lethal i.n. challenge with 1×10^4^ PFU of HSV-1, two-day old pups were administered 10 μg of mAb by i.p. injection. **A-C**. Survival of C57BL/6J pups receiving neutralizing mAb HSV8 or HSV8 LALA PG (**A**), non-neutralizing mAb CH42 AAA or CH42 NA (**B**), or isotype control mAb (**C**). Number of mice in each condition and statistical significance as compared to isotype control determined by the Log-rank (Mantel-Cox) test (*p<0.05, **p < 0.01, ***p<0.001, ****p < 0.0001) are reported in inset. Statistical significance between HSV8 and HSV8 LALA PG or CH42 AAA and CH42 NA are reported in the above legend as determined by the Log-rank (Mantel-Cox) test (**p < 0.01, ***p < 0.001, ****p < 0.0001). **D-F.** Viral titers were determined 5 days post infection (DPI). Data are shown as viral burden in perfused organs from surviving pups following 10 µg mAb treatment on DPI 0. Statistical significance was determined by 2-way ANOVA with Bonferroni’s test for multiple comparisons (**p < 0.01, ***p < 0.001, ****p < 0.0001). Geometric mean of the viral burden in organ type per treatment group is displayed. In legend n = number of pups included in viral titer of the total number of pups treated with mAb to account for pups who died prior to the time point of organ collection.

As an additional metric to explore the relative contributions of neutralization and effector functions in mediating protection, we assessed viral titers in various organs following 10 µg mAb treatment (**Figure 5 D-F)**. At 5 days post infection, both HSV8 and CH42 AAA significantly reduced viral burden in the brain, trigeminal ganglia (TG), and visceral organs (liver, spleen, and lungs) as compared to the isotype control mAb. HSV8 LALA PG, however, only significantly reduced viral burden in the brain as compared to isotype control (**Table S2**). The viral burden in pups treated with CH42 NA was indistinguishable from pups given an isotype control mAb. While not statistically significant, pups treated with HSV8 had lower viral burden as compared to pups given HSV8 LALA PG, consistent with survival data in indicating a contribution of effector functions in mediating protection **(Figure 5D)**. Further evidence for the role of effector functions in mediating protection was observed in the differences in viral burden in pups treated with CH42 AAA and CH42 NA. Pups treated with CH42 AAA had statistically significantly lower viral burden in the brain, TG, spleen, and lungs as compared to mice given CH42 NA **(Figure 5E)**. Of note, some pups given HSV8 LALA PG, CH42 NA, or the isotype control died prior to day 5 post infection, while no pups given HSV8 or CH42 AAA died prior to organ collection. Taken together, these data support a role for effector functions in protecting mice from HSV-1 mediated mortality and viral burden in the nervous system and viral dissemination.

### HSV-specific mAbs require effector functions for control of viral replication

To determine whether effector functions contribute to viral clearance, mouse pups were infected in a non-lethal challenge model utilizing a luciferase-producing recombinant HSV-1^35^, allowing real-time imaging of *in vivo* viral replication. Pups were challenged with HSV-1 17syn+dLux i.n., and were treated the following day with 10 µg of HSV8, HSV8 LALA PG, CH42 AAA, CH42 NA or an isotype control mAb delivered i.p.. Consistent with the results of survival and viral load experiments, mice that received a 10 μg dose of either HSV8 LALA PG or CH42 NA exhibited significantly greater levels of viral replication as measured by bioluminescence than mice treated with HSV8 or CH42 AAA starting at day 4 post infection (**Figure 6A-B**). Bioluminescence in Fc KO mAb-treated mice persisted for significantly longer than in those that received mAbs with intact effector functions and was comparable to the animals that received the IgG control mAb (**Figure 6A-B**). Statistically significant differences in bioluminescence were observed in animals treated with Fc KO versus Fc-functional mAbs (**Figure 6C**). Given the equivalent neutralization profiles of HSV8 and HSV8 LALA PG, differences in viral replication and dissemination must be attributable to the lack of effector functions in the LALA PG variant. Moreover, CH42 AAA, which does not neutralize, cleared virus significantly faster than its Fc KO counterpart. These results extend observations from the lethal challenge model and demonstrate that effector functions contribute to control of HSV-1 replication.

**Figure 6:**
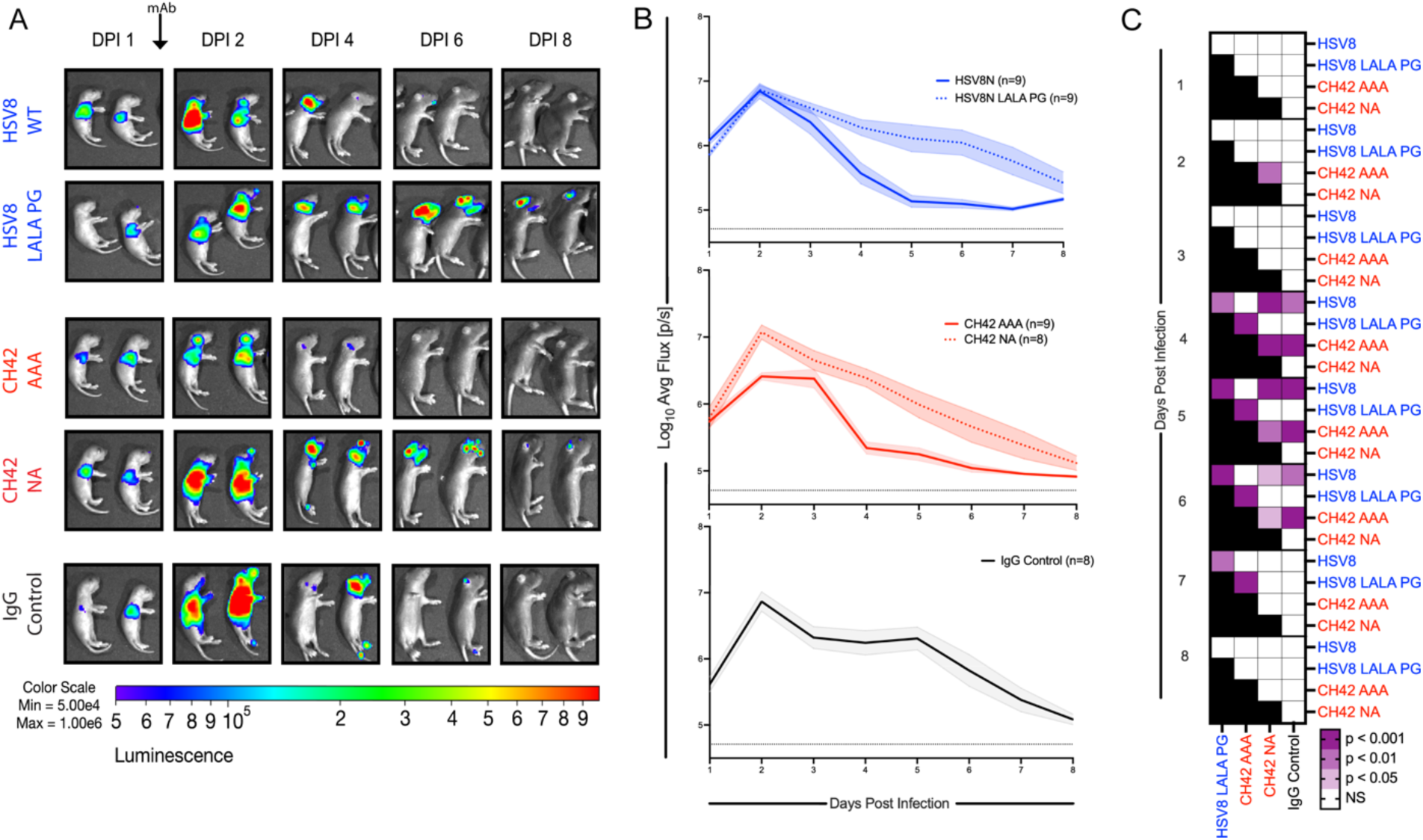
Effector functions accelerate control of viral replication after non-lethal HSV-1 challenge. One day post i.n. infection with a luciferase-expressing HSV-1, two-day-old pups were administered 10 µg of mAb i.p. and viral replication, as represented by bioluminescence, was quantified daily. **A**. Representative bioluminescence images of viral infection and replication following mAb treatment are presented for the same two pups over time. **B**. Quantification of virally-derived bioluminescence over time for HSV8 and HSV8 LALA PG (top), CH42 AAA and CH42 NA (middle), and isotype control (bottom). Lines and shaded regions represent the mean luminescence and standard error of the mean across pups (number listed in inset). **C**. Heatmap depicting statistical significance (2-way ANOVA with Tukey’s test for multiple comparisons) between groups treated with indicated mAbs over time after infection.

### Antibody functions contributing to protection differ between HSV serotypes

Since nHSV is caused by both HSV-1 and HSV-2, we next sought to examine whether the mechanism and patterns of protection were equivalent for both viruses. To test mechanism of protection against HSV-2, two-day old C57BL/6J mouse pups were treated with 40 µg of HSV-specific mAb or isotype control and then challenged with 300 PFU of HSV-2 strain G^36^. In contrast to HSV-1, and despite differences in neutralizing activities, both HSV8 and CH42 AAA provided equivalent protection against lethal challenge with HSV-2 (**Figure 7A,B**). Moreover, the Fc mutations in HSV8 LALA PG and CH42 NA completely ablated their protective activities, rendering them equivalent to the isotype control mAb (**Figure 7A-C**). These results demonstrate that Fc-mediated effector functions, and not viral neutralization, are essential for protection against HSV-2 infection, exposing a dichotomy between viral subtypes. Together, these data demonstrate that optimal antibody-mediated protection against HSV-1 in neonates is achieved by both neutralization and effector functions. In contrast, for protection against HSV-2, effector functions alone are sufficient.

**Figure 7:**
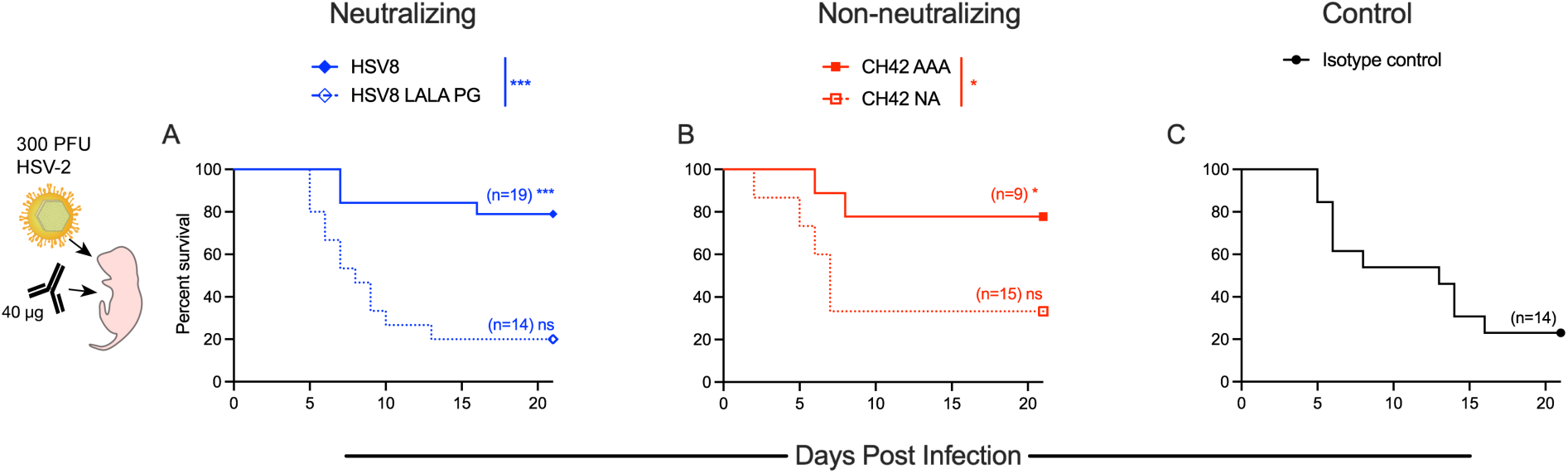
Antibody functions contributing to protection differ between HSV serotypes. Immediately before lethal i.n. challenge with 300 PFU of HSV-2, two-day old pups were administered 40 μg of mAb by i.p. injection. **A-C**. Survival of C57BL/6J pups receiving neutralizing mAb HSV8 or HSV8 LALAPG (**A**), non-neutralizing mAb CH42 AAA or CH42 NA (**B**), or isotype control mAb (**C**). Number of mice in each condition and statistical significance as compared to isotype control determined by the Log-rank (Mantel-Cox) test (*p<0.05, ***p<0.001) are reported in inset. Statistical significance between HSV8 and HSV8 LALA PG, or CH42 AAA and CH42 NA are reported in the above legend as determined by the Log-rank (Mantel-Cox) test (*p<0.05, ***p<0.001).

## Discussion

Understanding the mechanism by which antibodies provide protection has the potential to contribute to the development of mAb-based prevention and therapy, as well as to inform vaccine design. In this study, nHSV clinical outcomes depended on mAb specificity, neutralization potency, effector functions, dose, and viral strain. Both neutralization and effector functions improved virological outcomes following HSV-1 challenge. At higher Ab doses, neutralizing mAbs afforded near-complete protection, whereas the non-neutralizing mAb afforded only moderate protection, which was Fc-dependent since CH42 NA failed to prevent mortality. In contrast, under the same mAb dose and challenge conditions, viral neutralization alone was unable to prevent significant mortality in FcyR-/- mice. Notably, pre-incubating the virus with neutralizing mAb prior to challenging FcyR-/- mice completely protected these mice from mortality. This apparent discrepancy in the protective contribution of Fc-dependent antibody functions observed with KO mAbs versus FcyR-/- mice could be attributed to, for example, residual mAb effector function, differences in antibody biodistribution, and the intrinsic susceptibility of FcyR-/- mice^20^, among other factors.

When mAb was present at low concentrations, effector functions were more protective than viral neutralization. In contrast, when present at high levels systemically, or when pre-incubation with virus before nasal challenge, neutralization activity was the principal mechanism of action. This functional shift suggests that antibody concentration and biodistribution are determinants of the dominant mechanism of protection. Our findings support the hypothesis that antibody-mediated protection against HSV-1 is driven primarily by neutralization at high doses, while at lower doses both neutralization and Fc-effector functions play a role, as has been previously hypothesized^34^. Evidence in support of this hypothesis has been seen for other viruses. At sub-neutralizing antibody doses, mAb effector functions can be associated with improved resistance to infection^37,38^, control of viremia^39,40^, and clearance of virions^41^ during SHIV infection in non-human primates. Additionally, optimal mAb-mediated protection against SARS-CoV-2 infection required effector functions in addition to viral neutralization, particularly when neutralization potency was compromised^42–44^. Although these viruses differ from HSV in their pathogenesis and immune evasion strategies, our data support the idea that mAb dose is a pivotal determinant of the mechanism of protection. Antibody dose, however, can also directly impact clinical outcome in terms of viral pathogenesis. Sub-neutralizing doses of antibodies against Dengue virus can lead to FcR-driven antibody-dependent enhancement of disease^45^, furthering the consideration of antibody dose as a determinant for mechanism of action.

Given the ability of HSV8 LALA PG to protect against HSV-1, its relative inability to protect against HSV-2 was unexpected. The inability of both CH42 NA and HSV8 LALA PG to protect against HSV-2 indicates that Fc-mediated effector functions and not viral neutralization drive mAb-mediated protection against this serotype. This result may explain in part the failures of human HSV-2 vaccine trials^46–48^. A subunit vaccine containing gD and gB that induced high titers of neutralizing antibodies but low titers of ADCC-inducing antibodies^49^ showed poor efficacy^50^, indicating that neutralizing activity was not sufficient for prevention of genital disease and transmission of HSV-2. Similarly, a later gD subunit vaccine candidate that induced robust neutralizing titers but little to no ADCC activity^51^ had 58% efficacy in preventing HSV-1 genital disease, but could not prevent HSV-2 genital disease^46^. In this trial, neutralization titers against HSV-2 did not correlate with protection and could not explain the lack of vaccine efficacy^52^. Overall, the lack of protection afforded by neutralization and the poor effector function of antibodies raised by these vaccine candidates are consistent with the hypothesis that protection against HSV-2 requires effector functions. In our study, protection against HSV-2-mediated mortality was independent of mAb neutralization potency in that CH42 AAA poorly neutralized HSV-2 and yet provided protection comparable to HSV8. Consistent with this result, a non-neutralizing but FcγR-activating mAb that targets gB mediated protection from HSV-2 *in vivo*^53^ .

The importance of antibody effector functions was also observed in bioluminescent imaging experiments that sought to quantify viral load. Effector functions played the largest role in contributing to viral control, as both HSV8 and CH42 AAA cleared the HSV-derived bioluminescence significantly faster than their KO equivalents. Moreover, the ability for an antibody to elicit effector functions also greatly contributed to the control of viral burden and viral dissemination. Pups that received HSV8 or CH42 AAA had lower viral burden in tissues of the nervous system and in visceral organs as compared to their functional KO counterparts. This reduction in viral burden indicates a role for effector functions in the control of viral spread. HSV8 LALA PG was also able to slightly reduce viral burden in the brain as compared to the isotype control, indicating that neutralization still contributes to protection. Together, these preclinical studies highlight the importance of investigating non-neutralizing antibody functions in mediating protection against HSV disease, particularly HSV-2.

While there are caveats to direct translation of observations from animal models to humans, prior studies provide a high degree of confidence as to which murine FcγR are engaged when introducing human IgG1 into a mouse^22,54^. The distribution of FcRs varies between human and murine innate immune cells, but the overall effector functions elicited by the differing cell types are conserved. ADCC activity elicited via human cells is generally a good predictor of murine ADCC (predominantly carried out by macrophages and PMNs)^55^. We focused on mAbs specific for gD and tested a limited number of viral strains in a single mouse strain background, and our results may or may not generalize across other target antigens, mAbs, viruses, or host genetic backgrounds. Other caveats include when and where mAbs initially encounter virus, particularly in the context of differing hosts. Given that humans show a spectrum of anatomical, physiological, and immunological profiles, and based on the data of this study, antibodies with broad functional activities are more likely to afford clinical efficacy. This idea is supported by clinical evidence: both neutralizing and ADCC Ab activity serve as biomarkers for protection of infants from disseminated HSV disease^10^. The inability of neutralizing activity to serve as a reliable biomarker of vaccine-mediated protection in adults, particularly for HSV-2, are also consistent with our results. Collectively, these data support the conclusion that polyfunctional mAbs able to elicit both neutralization and effector functions are the best candidates for therapeutic and prophylactic translation. Expanding the focus of vaccine research and development to include activities beyond viral neutralization has the potential to accelerate the quest for interventions to reduce the global burden of HSV infection.

## Acknowledgments

We would like to thank all members of the Ackerman and Leib laboratories for their invaluable help with critical discussion and scientific advice. We thank United BioPharma for providing UB-621 and ZabBio for providing HSV8. The authors acknowledge the following Shared Resources facilities: Irradiation, Pre-clinical Imaging and Microscopy Resource (IPIMSR) at the Norris Cotton Cancer Center at Dartmouth with NCI Cancer Center Support Grant 5P30 CA023108-37. These studies were partially supported by National Institutes of Health grants, NEI 5R01EY009083-28 to DAL, NIAID grants 5P01AI098681-08 and 5R21AI147714-02 to DAL, U19AI145825 to MEA, R01AI176646 to DAL and MEA, and T32AI007363 to MDS.

## Author Contributions

IMB, MDS, DAL, and MEA conceptualized the study. IMB, MDS, NSK, and CRG performed experiments. DAL and MEA obtained funding and supervised research. IMB and MDS drafted manuscript and generated figures. IMB, MDS, DAL, and MEA finalized manuscript and all other authors have read and edited.

## Declaration of Interests

IMB, DAL, and MEA report a patent WO2020077119A1 for mAbs used in this manuscript as a method for the treatment for nHSV infections.

## Methods

### RESOURCE AVAILABILITY

#### Lead Contact

Further information and requests for reagents and resources should be directed to and will be fulfilled by the lead contact Dr. Margaret E. Ackerman

#### Materials Availability

Antibodies, cell lines, and plasmids generated for this study may be requested with a material transfer agreement.

#### Data and code availability

All data reported in this paper will be shared by the lead contact upon reasonable request. This paper does not report original code.

Any additional information required to reanalyze the data reported in this paper is available from the lead contact upon request.

### EXPERIMENTAL MODEL AND STUDY PARTICIPANT DETAILS

#### Cell Lines

Vero Cells (CCL-81) were purchased from American Type Culture Collection (ATCC) and were maintained in Dulbecco’s Modified Eagle’s Medium (DMEM) containing 5% fetal bovine serum (FBS) and 1% penicillin/streptomycin at 37°C with 5% CO_2_. HEK293Ts were purchased from ATCC and maintained in DMEM with 10% FBS at 37°C and 5% CO_2_. The human monocytic cell line, THP-1, was purchased from ATCC and maintained in RPMI-1640 supplemented with 10% FBS and 55µM beta-mercaptoethanol at 37°C with 5% CO_2_. EXPI293Fs were purchased from ThermoFisher and were maintained in Expi293F Media (Thermo Fisher). Cells were grown in a Thermo Scientific reach-in CO_2_ incubator at 37°C with 8% CO_2_ on an innOva 2300 platform shaker at 125 RPM. Jurkat-Lucia NFAT CD16 cells were purchased from Invivogen and grown in RPMI-1640 supplemented with 10% fetal bovine serum, 1 mM sodium pyruvate, 1x non-essential amino acids, 1x penicillin/streptomycin, 100 µg/mL Normocin, 100 µg/mL Zeocin, and 10 µg/mL Blasticidin.

#### Animals

Naïve male and female C57BL/6J (RRID: IMSR_JAX:000664) were either purchased from The Jackson Laboratories or bred in animal facilities at Dartmouth College in accordance with institutional animal care and use committee protocols (Dartmouth College IACUC 2151). C57BL/6J mice were bred according to IACUC protocols and 2-day-old offspring of both sexes were then used in challenge studies. Naïve male and female B6.129P2-Fcer1gtm1Rav N12 (FcyR-/-) (model: 583) were purchased from Taconic Labs. FcyR-/- mice were bred in accordance with IACUC protocols and 2-day-old offspring of both sexes were used in challenge studies.

### METHOD DETAILS

#### Mouse procedures and viral challenge

C57BL/6J (B6) mice were purchased from The Jackson Laboratory. FcγR-/- mice (B6.129P2-Fcer1gtm1Rav N12) were purchased from Taconic Labs^33^. Administration of mAbs was via the peritoneal route with a 25 μL Hamilton syringe in a 20 μL volume under 1% isoflurane anesthesia. The wild-type viral strains used in this study were HSV-1 17syn+ ^56^, HSV-2 G (kindly provided by Dr. David Knipe)^36^. The bioluminescent luciferase-expressing recombinant virus HSV-1 17syn+/Dlux was constructed as previously described^35^. Viral stocks were prepared using Vero cells as previously described^57,58^. Newborn pups were infected i.n. on day 2 postpartum with indicated amounts of HSV in a volume of 5-10 μl under 1% isoflurane anesthesia. Pups were then monitored for survival, imaging, or viral burden analysis. For survival studies, pups were challenged with 1×10^4^ plaque forming units (PFU) of HSV-1 (Strain 17), and 3×10^2^ PFU of HSV-2 (Strain G), as indicated. Endpoints for survival studies were defined as excessive morbidity (hunching, spasms, or paralysis) and/or >10% weight loss (**Figures S3-5**). For bioluminescent detection, pups were injected i.p. with 20 µl of 15 mg/mL D-luciferin potassium salt (Gold Biotechnology), placed in isoflurane chamber, and moved into a Xenogen IVIS-200 with a warmed stage and continuous isoflurane. Pups were typically imaged beginning at 1 day post-infection and serially imaged every day until 8 days post-infection to monitor bioluminescence. For viral titers of organs, tissues were harvested 5 days post infection following cardiac perfusion with at least 5 mL of ice-cold PBS. All tissues were collected in 1.7 mL tubes containing ∼100 µL of 1mm sterile glass beads and 1 mL of DMEM containing 5% fetal bovine serum (FBS) and 1% penicillin/streptomycin. Tissue homogenates were prepared via mechanical disruption using a Mini-Beadbeater-8 (BioSpec Products). Organ titers were measured via plaque assay on Vero cells.

#### Monoclonal antibodies

CH42^14^ AAA plasmids were kindly provided by Dr. Anthony Moody (Duke University). When expressed *in vitro*, CH42 contained the Fc mutation known as AAA (S298A/E333A/K334A), which enhances antibody dependent cellular cytotoxicity^27^. The variable heavy chain sequence of CH42 was subcloned into a plasmid coded with an IgG1 heavy chain backbone containing the N297A ^29^ mutation via QuikChange Site Directed Mutagenesis kit (Agilent). Antibodies were expressed through co-transfection of heavy and light chain plasmids in Expi293 HEK cells (Thermo Fisher) according to the manufacturer’s instructions. Seven days after transfection, cultures were spun at 3000 x g for 30 minutes to pellet the cells, and supernatants were filtered (0.22 μm). IgG was affinity purified using a custom packed 5 mL protein A column with a retention time of 1 minute (ie. 5 mL/min) and eluted with 100 mM glycine pH 3, which was immediately neutralized with 1 M Tris buffer pH 8. Eluate was then concentrated to 2.5 mL for size exclusion chromatography on a HiPrep Sephacryl S-200 HR column using an AktaPure FPLC at a flow rate of 1 mL/min of sterile PBS. Fractions containing monomeric IgG were pooled and concentrated using spin columns (Amicon UFC903024) to approximately 2 mg/mL of protein and either used within a week or aliquoted and frozen at - 80°C for later use. HSV8 mAb was kindly provided by ZabBio and Kentucky Bioprocessing, and a clinical grade antibody preparation of UB-621 was kindly provided by United BioPharma.

#### Measurement of antibody binding to mouse and human Fc Receptors

Recombinant gD antigen ^59^, kindly provided by Dr. Gary Cohen (UPenn), was coupled to MagPlex beads (Luminex) as previously described ^60^. gD mAbs were serially diluted in 1x PBS with 0.1% bovine serum albumin (BSA) and 0.05% Tween-20 and incubated with antigen-coupled beads overnight at 4°C with constant shaking. Beads were washed before being incubated with recombinant biotinylated human Fc receptors^61^ (Duke Human Vaccine Institute) or mouse Fc receptors (Sino Biologics) that were tetramerized with streptavidin-PE for 1 hour. The beads were washed and analyzed on the xMap system. The median fluorescence intensity of at least 10 beads/region was recorded. An isotype control antibody and a buffer only control were used to determine antigen-specific binding and assay background signal. Area under the curve was calculated using Prism 9 (GraphPad).

#### Viral Neutralization

Serially diluted mAb and 50 PFU of HSV-1 st17 or HSV-2 G were incubated for 1 hour at 37°C before being added to confluent Vero cells grown in 6 well plates. Immune complexes were incubated with Vero cell monolayers for 1 hour at 37°C with 5% CO_2_ with shaking every 15 minutes. Methylcellulose overlay was added to the wells after the hour incubation. Plates were incubated for 48 (HSV-1) or 72 (HSV-2) hours at 37°C with 5% CO_2_. Methylcellulose overlay was removed, Vero cells were fixed with 1:1 ethanol:methanol before being stained with 12% Giemsa overnight. Stain was removed and plaques were counted on a light box. Virus neutralization (%) was calculated as [(# of plaques in virus only - # of plaques counted at mAb dilution)/# of plaques in virus only well] ×100.

#### Antigen Binding ELISA

The ability for the HSV-specific mAbs to bind to gD was evaluated via an ELISA. Briefly, the wells of a high-binding 96 well plate were coated with 1 µg/mL gD in sodium bicarbonate buffer pH 9.4 and incubated overnight at 4°C. The plates were washed 5x with 1x PBS, 0.1% BSA, 0.05% Tween-20 and blocked with 1x PBS with 2.5% BSA overnight at 4°C. The plates were washed 5x. Antibodies were serially diluted in 1x PBS with 0.1% BSA over a seven point two-fold dilution curve (10.66 nM – 0.16 nM), added to the plates, and incubated at room temperature for 1 hour. The wells were washed 5x and incubated with 100 µl/well with an HRP-conjugated anti-human IgG Fc antibody (1:10000 dilution, Invitrogen) for 1 hour. Wells were washed a final time before being incubated with 100 µL/well 1-step Ultra TMB (Invitrogen) for 5 minutes. The reaction was halted with 100 µL/well 1N H_2_SO_4_. The plate was read at 450 nm on a SpectraMax Paradigm Plate Reader (Molecular Devices). Buffer only wells were used as a control and the assay was performed in technical replicate.

#### Antibody-dependent cellular cytotoxicity (ADCC)

A CD16 activation reporter assay was performed as previously described^62^. Briefly, the wells of a high-binding 96 well plate were coated with 1 µg/mL recombinant gD protein in PBS and incubated overnight at 4°C. The plate was washed 3x with 1x PBS with 0.01% Tween20 and blocked at room temperature with 1x PBS with 2.5% BSA for 1 hour. Antibodies were serially diluted in growth medium and added to the washed plate with 100,000 Jurkat Lucia NFAT CD16 cells/well (Invivogen). Antibodies and cells were incubated for 24 hours at 37°C with 5% CO_2_. A 25 µL volume of the cell supernatant was removed and added to a new, opaque white 96 well plate. A 75 µL volume of the QuantiLuc (Invivogen) substrate was added to the supernatant and luminescence was immediately read on SpectraMax Paradigm plate reader (Molecular Devices) using a 1 second integration time. A kinetic read time of 0, 2.5 and 5 minutes was performed, and the reported values are the averages of the three reads. Buffer only wells were used as negative controls and a cell stimulation cocktail with 2 µg/mL ionomycin was used as a positive control. The assay was performed in technical replicate.

#### Antibody-dependent cellular phagocytosis (ADCP)

Antibody-dependent cellular phagocytosis was performed as previously described ^63^ with slight modifications. Briefly, goat-anti human IgG F(ab’)2 (Invitrogen) was covalently coupled to yellow-green carboxylate beads (Thermofisher). Antibodies were diluted in culture medium to a starting concentration of 133 nM and serially diluted 4-fold 7 times. Diluted mAbs were incubated with anti-human IgG beads for 2 hours at 37°C to form immune complexes. THP-1 (ATCC) cells (25,000/well) were added to the immune complexes and incubated at 37°C for 4 hours. Cells were washed 2x with cold 1x PBS prior to being fixed with 4% paraformaldehyde. The cells were analyzed on a NovoCyte Advanteon flow cytometer (Agilent) (**Figure S2C)**. A phagocytosis score was calculated as the (percentage of FITC+ cells) x (the geometric mean fluorescence intensity (gMFI) of the FITC+ cells)/100,000. Buffer only wells were used as negative controls and the assay was performed in technical replicate with two biological replicates.

#### Engineering HEK293Ts expressing HSV-1 gD as a surface antigen

The gD gene was PCR amplified from HSV-1 strain 17 DNA. The gene was cloned into pLenti-DsRed-IRES-EGFP vector (Addgene plasmid number 92194)^64^ by restriction digestion using Afe1 and BamH1 (New England BioLabs (NEB)). Restriction digestion was followed by ligation using T4 DNA ligase (NEB). The ligated PCR product was transformed into NEB® Stable Competent *E. coli* (High Efficiency). The gene insertion into the vector was confirmed by using restriction digestion by SgrA1 (NEB) and plasmid sequencing (Azenta LifeSciences). The sequence confirmed plasmid (transfer plasmid) and packaging vector (VSVG, PSPAX2) were used at concentrations of 6 μg, 0.6 μg and 5.4 μg to transfect HEK293T cells at 60% confluency in a T150 flask. Transfer plasmid and packaging vector were mixed with Opti-MEM (ThermoFisher Scientific). In a separate tube, Opti-MEM and 109.38 μg Polyethylenimine (PEI) was added. The DNA:Opti-MEM and PEI:Opti-MEM mixtures were combined and incubated together for 15 min at room temperature prior to being added to the HEK-293Ts. Media was replenished the next day (day 1). On day 2, the viral supernatant was collected and Lenti-X™ GoStix™ Plus (Takara) was used to test presence of lentiviral p24. The viral supernatant was filtered using 0.45-micron filter and aliquots were stored at -80 °C.

Adherent HEK293T cells were trypsinized and 500,000 cells were mixed in 1 mL of thawed viral supernatant, to which 0.8 μg of polybrene (Santa Cruz Biotechnology) was added. The mixture was incubated in a 6 well plate at 37 °C, 5% CO_2_. On the next day, old media was removed and was replaced with 2 mL fresh media. At 4 days post transduction, GFP positive cells were sorted using cell sorter (Sony Biotechnology, MA900) using a 100-micron sorting chip (Sony Biotechnology) and cultured in media containing 1X penicillin/streptomycin. Non-transfected HEK293T cells were used as a negative control to set the sort gates (**Figure S2)**.

#### Measurement of binding of antibody to HEK293Ts expressing HSV-1 gD as a surface antigen

HEK293T cells expressing HSV-1 gD as a surface antigen and non-transfected HEK293T cells (control) were washed twice with PBS. Cells (200,000/well) were added to a 96 well V bottom plate (USA Scientific). gD-specific antibodies were diluted to 20 µg/mL and serially diluted four-fold in PBS + 1% BSA before being added to the cells. After a 1-hour incubation on ice, the cells were washed twice with PBS + 1% BSA and stained with 10 μg/mL Alexa Fluor™ 647 Goat anti-Human IgG (H+L) Cross-Adsorbed Secondary Antibody (ThermoFisher Scientific) diluted in PBS with 1% BSA. After a 30 min incubation in the dark, cells were washed twice with PBS + 1% BSA and were resuspended in 100 μL of PBS prior to fixation with 4% paraformaldehyde. The antibody binding was measured by checking signal intensity of Alexa Fluor 647 using a MACSQuant Analyzer (Miltenyi) (**Figure S2B**). The experiment had two biological replicates. The data was analysed using FlowJo version 10.8.2.

#### Antibody Dependent Complement Deposition

HEK293T cells expressing HSV gD as surface antigen and non-transfected HEK293T cells (control) were washed twice with PBS. Cells (200,000/well) were added to a 96 well V bottom plate (USA Scientific). Antibodies were diluted to 20 µg/mL and serially diluted four-fold in PBS + 1% BSA before being added to the cells. After 45 min, the cells were washed with PBS + 1% BSA, followed by a wash with Gelatin Veronal Buffer (GVB++) (Complement Technology Inc). Low-tox Guinea Pig complement (Cedarlane) was reconstituted in 1 mL cold distilled water, a 500 µL volume of which was added to 9.5 mL GVB++. Diluted guinea pig complement (100 µL) was then added to each well prior to incubation with orbital shaking for 1 hour at 37 °C, with 5% CO_2_. The cells were then washed with PBS + 1% BSA prior to staining with 100 µL of 1 µg/mL biotinylated goat anti-guinea pig C3 antibody (ICL labs) at room temperature for 1 hour. The cells were washed twice with PBS + 1% BSA prior to addition of 100 µL of 1 µg/mL Streptavidin-APC (ThermoFisher) and incubation for 1 hour at room temperature. After the incubation, the cells were washed twice and resuspended in PBS + 1% BSA. Antibody-dependent activation of complement protein C3 was measured using a MACSQuant Analyzer (Miltenyi) quantifying the mean fluorescence intensity of APC (**Figure S2A**). The assay was performed with two biological replicates. Heat-inactivated guinea pig complement was used as a control. For heat inactivation, the serum was heated at 58°C for 30 min. VRC01 antibody was used as a negative control. The data was analysed using FlowJo version 10.8.2.

#### CD16 activation assay (ADCC)

HEK293T cells expressing HSV gD as surface antigen and non-transfected HEK293T cells were washed 2x with PBS before being added to a V bottom plate (USA Scientific) (200,000 cells/well). Into the same plate, 100,000 cells/well of Jurkat Lucia NFAT CD16 cells (Invivogen) were added, along with 180 µL of assay media (RPMI 1640 + 10% FBS + 1mM sodium pyruvate + non-essential amino acids + penicillin/streptomycin) and 20 µL of diluted gD-specific antibodies (in PBS + 1% BSA). The plate was incubated overnight at 37°C, 5% CO_2_. After overnight incubation, the cells were centrifuged and 25 µL of supernatant was drawn from each well and transferred into 96-well white walled clear bottom polystyrene plate (Costar) and mixed with 75 µL of reconstituted QUANTI-Luc™ reagent (InvivoGen). Luminescence was immediately read on a SpectraMax Paradigm Plate reader (Molecular Devices) using 1s integration time. Kinetic reads at 0 min, 2.5 min and 5 min were measured, and the mean reading was noted. Cell Simulation Cocktail (eBioscience) was used as positive control. VRC01 was used as negative control. The assay was performed with two biological replicates.

#### Study Approval

Procedures were performed in accordance with Dartmouth’s Center for Comparative Medicine and Research policies and following approval by the institutional animal care and use committee.

#### Statistical Analysis

Prism 9 (GraphPad) software was used for statistical tests. For survival studies, HSV-specific mAbs were compared to isotype controls using the Log-rank Mantel-Cox test to determine *p* values. HSV-specific Fc-competent and KO mAbs were also compared to each other using the Log-rank Mantel-Cox test to determine *p* values. For imaging studies, groups and time points were compared to each other via 2-way ANOVA, with Tukey’s test for multiple comparisons to determine *p* values. For viral burden analysis, mAbs were compared to each other within each organ group via an ordinary 2-way ANOVA with Bonferoni’s test for multiple comparison. Within each organ, HSV-specific mAbs were compared to the isotype control mAb via a 2-way ANOVA with Dunnet’s test for multiple comparisons.

