## Supplementary Materials for "Antibody effector functions are required for broad and potent protection of neonates from herpes simplex virus infection"

| **Supplemental Figures** | |
|---|---|
| Supplemental Figure 1 | HSV gD mAbs display altered binding profiles to human and mouse Fcγ Receptors. |
| Supplemental Figure 2 | Representative Flow Cytometry Plots. |
| Supplemental Figure 3 | Percent weight change of mouse pups receiving 40 µg of HSV-specific mAbs. |
| Supplemental Figure 4 | Percent weight change of mouse pups receiving Immune Complexed mAb:Virus. |
| Supplemental Figure 5 | Percent weight change of pups receiving HSV-specific mAbs following HSV infection. |
| **Supplemental Tables** | |
| Supplemental Table 1 | Fc variants used in study and their properties. |
| Supplemental Table 2 | Adjusted P-values for comparisons between treated groups and viral titers in organs. |

Figure S1

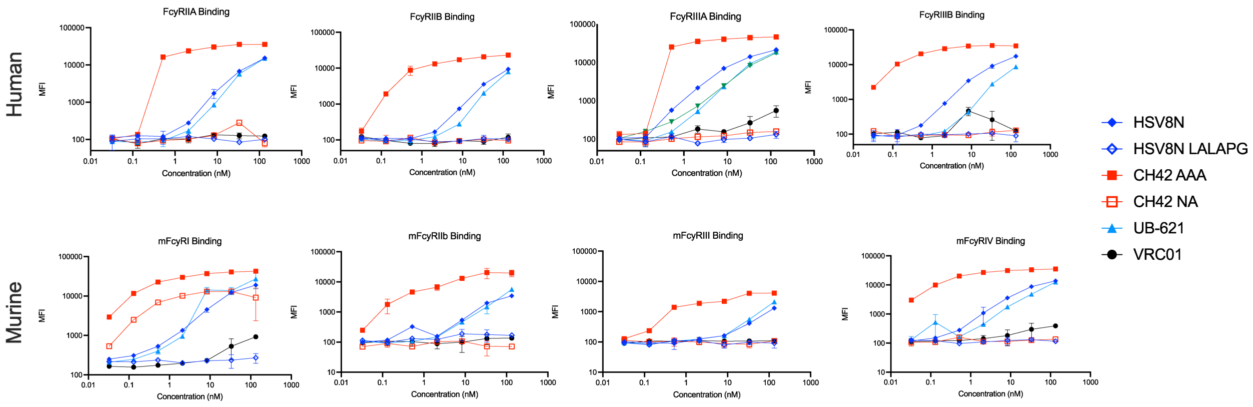


**Figure S1 (related to figure 1). HSV gD mAbs display altered binding profiles to human and mouse Fcγ Receptors.** Raw FcR binding profiles for HSV-specifc mAbs bound to gD and detected with human (top) or mouse (bottom) Fc receptors tetramerized with streptavidin-PE. Error bars represent standard deviation from the mean. MFI – median fluorescence intensity.

Figure S2

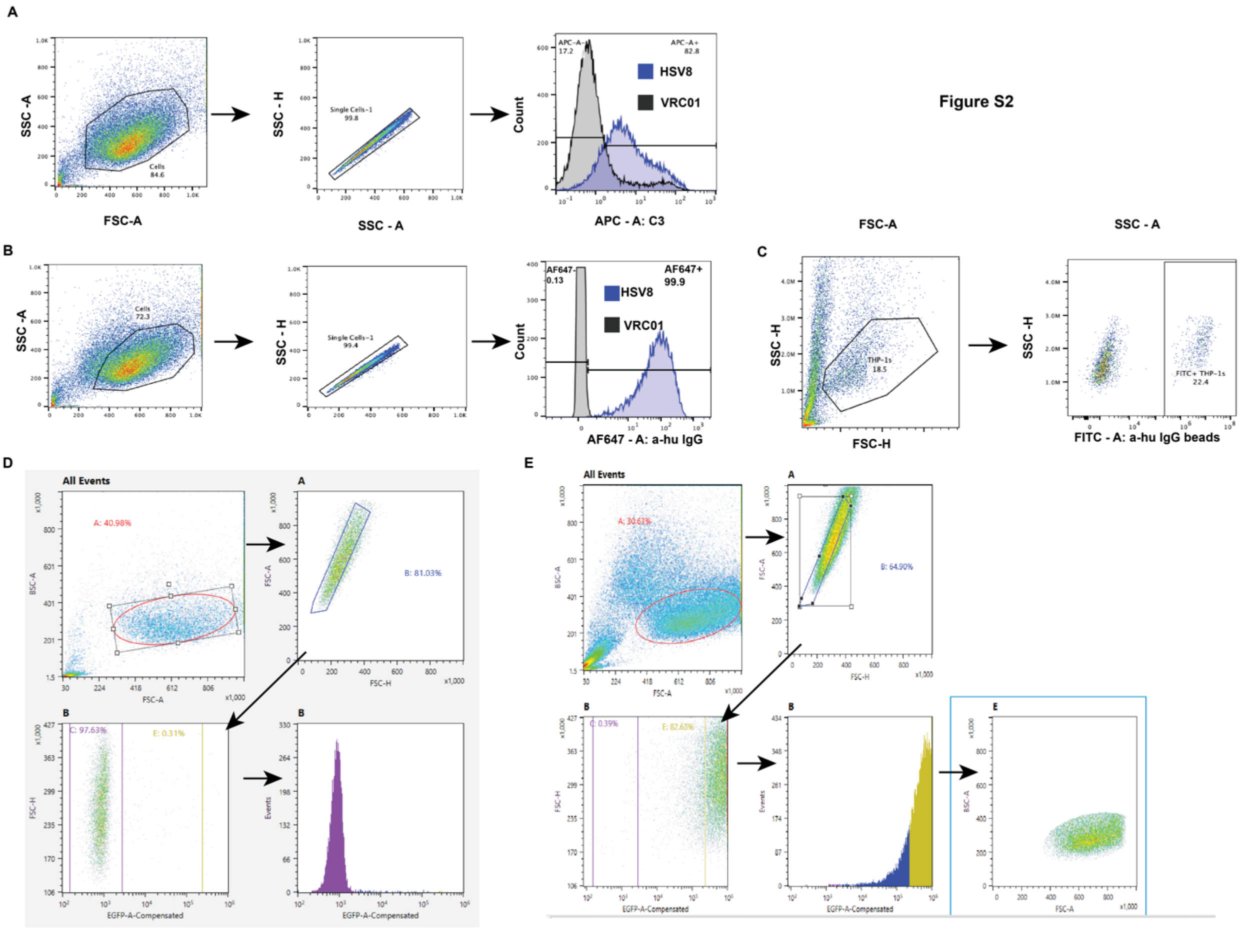


**Figure S2 (related to figure 2). Representative Flow Cytometry Plots.** Representative flow cytometry plots and gating strategies are shown for **A.** antibody-dependent complement deposition (ADCD), **B.** antibody binding to HSV-gD expressed on HEK293Ts, and **C.** antibody-dependent cellular phagocytosis (ADCP). **D.** Gating strategy for the sorting of non-transfected HEK293Ts as a control. **E.** Gating strategy for the sorting of HEK293Ts expressing HSV-1 gD on the surface using GFP as a selection marker.

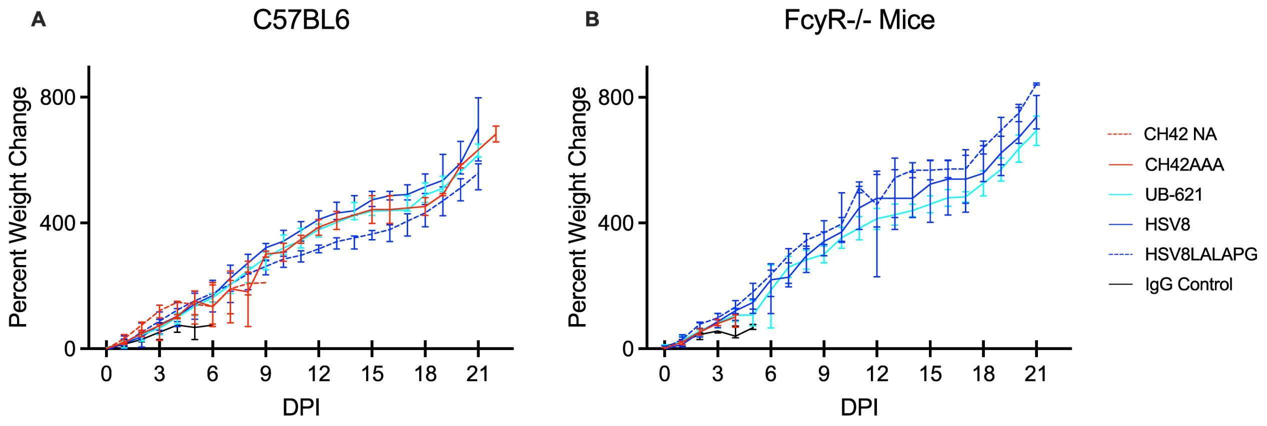


**Figure S3 (related to figure 3). Percent weight change of mouse pups receiving 40 µg of HSV-specific mAbs**. Immediately before lethal intranasal challenge with 1e4 plaque forming units (PFU) of HSV-1, two-day old pups were administered 40 µg of mAb by intraperitoneal (IP) injection. **A**. Weight change of C57BL/6J mice receiving mAb and following intranasal HSV-1 challenge. **B.** Weight change of FcyR-/- mice receiving mAb following intranasal HSV-1 challenge. DPI – Days Post Infection. The median baseline corrected weight of each treatment group is plotted. Error bars represent the 95% confidence interval.

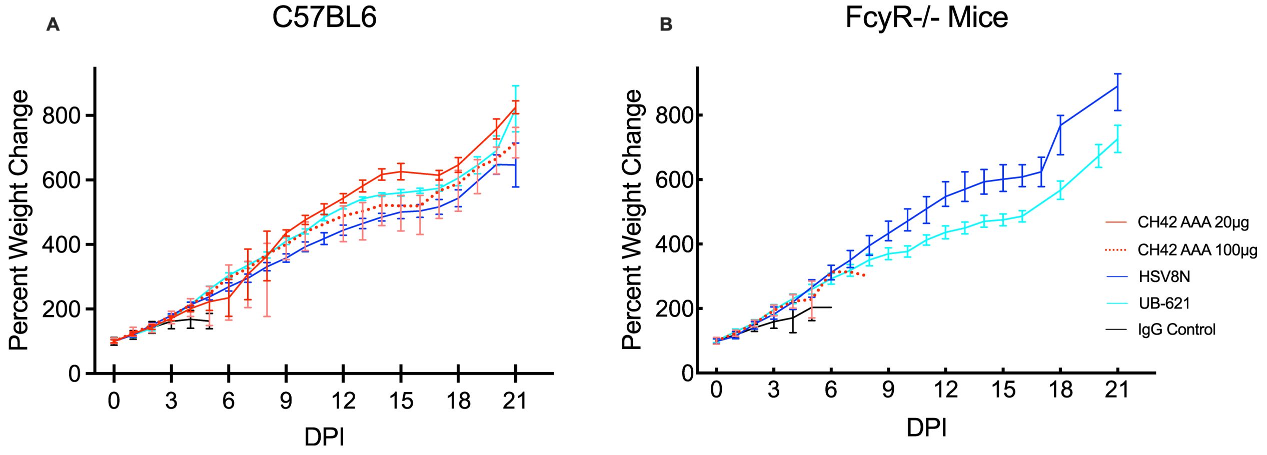


**Figure S4 (related to figure 4). Percent weight change of mouse pups receiving Immune Complexed mAb:Virus.** One hour before intranasal challenge of two-day old pups, immune complexes were formed by incubation of 1e4 plaque forming units (PFU) of HSV-1 with mAb at 37˚C (20 mg unless otherwise noted. **A**. Weight change of C57BL/6J mice receiving immune complexed HSV-1. **B**. Weight change of FcyR-/- mice receiving immune complexed HSV-1. DPI – Days Post Infection. The median baseline corrected weight of each treatment group is plotted. Error bars represent the 95% confidence interval.

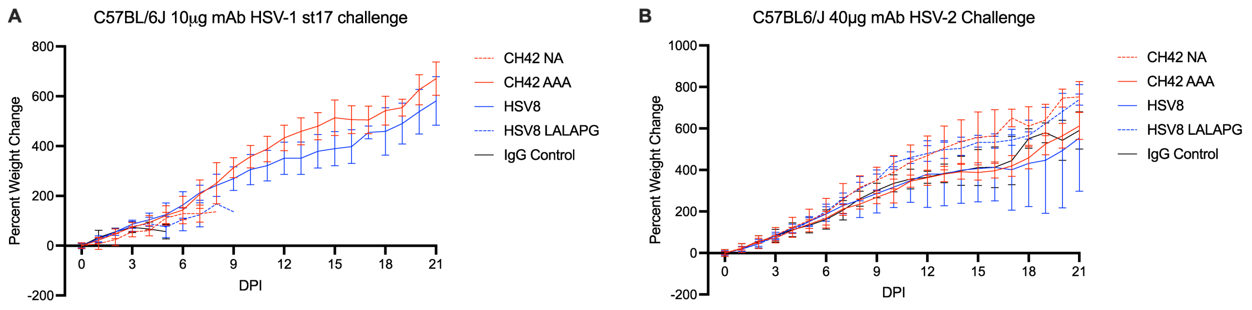


**Figure S5 (related to figures 5 and 7). Percent weight change of pups receiving HSV-specific mAbs following HSV infection A**) Immediately before lethal intranasal challenge with 1e4 plaque forming units (PFU) of HSV-1, two-day old pups were administered 10 ug of mAb by intraperitoneal (IP) injection. Pups were weighed daily and percent weight change for each group is reported in A. **B**) Immediately before lethal intranasal challenge with 300 plaque forming units (PFU) of HSV-2, two-day old pups were administered 40 ug of mAb by intraperitoneal (IP) injection. Pups were weighed daily and percent weight change from baseline is reported in B. DPI – Days Post Infection. The median baseline corrected weight of each treatment group is plotted. Error bars represent the 95% confidence interval.

**Table S1**

| Fc variant name | Function | | | | References |
|---|---|---|---|---|---|
| | ADCC | ADCP | ADCD | 1/2 life (Hu) | |
| WT (Afucosylated) | ↑↑ | ←→ | ←→ | ←→ | |
| AAA | ↑↑ | ←→ | ←→ | ←→ | 27 |
| LALA PG | ↓↓ | ↓↓ | ↓↓ | ←→ | 29 |
| NA | ↓↓ | ↓↓ | ↓↓ | ←→ | 30 |

Enhanced: ↑↑, unchanged: ←→, slightly diminished: ↓, diminished: ↓↓, non-functional: ----,

**Table S1 (related to figure 1) Fc variants used in study and their properties.**

**Table S2**

| Comparison | Adjusted p-value | Summary |
|---|---|---|
| **HSV8 (n=7) vs HSV8 LALA PG (n=7)** | | |
| Brain | >0.9999 | ns |
| TG | >0.9999 | ns |
| Liver | >0.9999 | ns |
| Spleen | 0.6821 | ns |
| Lungs | 0.5684 | ns |
| **CH42 AAA (n=9) vs CH42 NA (n=5)** | | |
| Brain | <0.0001 | **** |
| TG | 0.0005 | *** |
| Liver | 0.1403 | ns |
| Spleen | 0.0083 | ** |
| Lungs | 0.0011 | ** |
| **Brain** | | |
| Isotype control (n = 9) vs. HSV8N (n=7) | 0.0024 | ** |
| Isotype control (n = 9) vs. HSV8 LALA PG (n=7) | 0.0061 | ** |
| Isotype control (n = 9) vs. CH42 AAA (n=9) | | |
| Isotype control (n = 9) vs. CH42 NA (n=5) | <0.0001 | **** |
| | >0.9999 | ns |
| **TG** | | |
| Isotype control (n = 9) vs. HSV8N (n=7) | 0.0031 | ** |
| Isotype control (n = 9) vs. HSV8 LALA PG (n=7) | 0.0993 | ns |
| Isotype control (n = 9) vs. CH42 AAA (n=9) | <0.0001 | **** |
| Isotype control (n = 9) vs. CH42 NA (n=5) | 0.9748 | ns |
| **Liver** | | |
| Isotype control (n = 9) vs. HSV8N (n=7) | 0.0329 | * |
| Isotype control (n = 9) vs. HSV8 LALA PG (n=7) | 0.3112 | ns |
| Isotype control (n = 9) vs. CH42 AAA (n=9) | <0.0001 | **** |
| Isotype control (n = 9) vs. CH42 NA (n=5) | 0.1732 | ns |
| **Spleen** | | |
| Isotype control (n = 9) vs. HSV8N (n=7) | 0.0011 | ** |
| Isotype control (n = 9) vs. HSV8 LALA PG (n=7) | 0.3381 | ns |
| Isotype control (n = 9) vs. CH42 AAA (n=9) | <0.0001 | **** |
| Isotype control (n = 9) vs. CH42 NA (n=5) | 0.6401 | ns |
| **Lungs** | | |
| Isotype control (n = 9) vs. HSV8N (n=7) | 0.0021 | ** |
| Isotype control (n = 9) vs. HSV8 LALA PG (n=7) | 0.5405 | ns |
| Isotype control (n = 9) vs. CH42 AAA (n=9) | <0.0001 | **** |
| Isotype control (n = 9) vs. CH42 NA (n=5) | 0.9251 | ns |

**Table S2 (related to figure 5) Adjusted P-values for comparisons between treated groups and viral titers in organs.** 2-way ANOVA with Bonferroni's test for multiple comparisons was used to determine statistical significance between viral titers in organs between mice given HSV-specific antibodies. The wild-type antibody was compared to the Fc KO mAb in each case. 2-way ANOVA with Dunnett's test for multiple comparisons was used to compare viral titers in organs of pups receiving HSV-specific antibodies to mice who received isotype control mAb.
